# The role of local and long-range stresses in cephalic furrow formation in the *Drosophila melanogaster* embryo

**DOI:** 10.1101/2024.12.30.630777

**Authors:** Redowan A. Niloy, Guo-Jie J. Gao, Michael C. Holcomb, Jeffrey H. Thomas, Jerzy Blawzdziewicz

**Affiliations:** Department of Mechanical Engineering, Texas Tech University, Lubbock, TX, U.S.A.; Department of Aerospace and Mechanical Engineering, University of Notre Dame, Notre Dame, IN, U.S.A.; Department of Mathematical and Systems Engineering, Shizuoka University, Hamamatsu, Japan; Department of Physics and Geosciences, Angelo State University, San Angelo, TX, U.S.A.; Department of Cell Biology and Biochemistry, Texas Tech University Health Sciences Center, Lubbock, TX, U.S.A.; Department of Physics and Astronomy, Texas Tech University, Lubbock, TX, U.S.A.

## Abstract

Cephalic furrow (CF) is a transient epithelial invagination that forms during early gastrulation in the *Drosophila melanogaster* embryo. The initial stage of cephalic furrow formation (CFF) involves a shortening of initiator cells, generation of a localized asymmetric protrusion inwards, and then subsequent descent of cells into the yolk sac area. We present an analysis of how local forces associated with cell-membrane tensions and cell pressures interact with the long-range tensile stress developing along the furrow to generate the invagination. We propose two numerical models which capture different aspects of CFF. First, we formulate a force-center model of CF to show how the spatiotemporal heterogeneity of initiator-cell activation observed *in vivo* is a result of tensile-stress-feedback-based intercellular coordination. We also argue that this kind of mechanical stress-based activation mechanism likely contributes to robustness of the overall process. Second, we use our multi-node lateral vertex model to analyze the mechanical dynamics of the anterior–posterior cross-section of CF. This approach allows us to quantify the balance between cortical membrane tension forces, cellular pressures, and the inward force produced by the tension along the curved apical surface of the embryo. Comparing our simulations to experimental images, we discuss the crucial and indispensable role of the tension-induced inward force, especially during the initial stages of CFF where the localized asymmetric protrusion is formed. We argue that without this inward force the initial descent of the initiator cells into the furrow would not be possible, and that at later stages the inward force provides redundancy to this process and thus aids CFF robustness.

## I. INTRODUCTION

Cell-level and tissue-scale mechanical forces and the associated transduction and feedback play a crucial role in developmental processes such as generation of embryonic architecture and organ formation [1–6]. Mechanical effects are present even in the early blastoderm [7]. Mechanical forces are also important in maintenance of healthy tissues such as bones [8], muscles, and blood vessels [9]. Moreover, mechanically controlled processes are implicated in disease development, e.g., in pathological tissue remodeling in wound healing [10–12] and in proliferation of cancer cells [13–17]. Thus, investigations of the role of mechanical forces and mechanical sensing in biological systems are essential, and this research field is rapidly expanding.

In embryonic development, mechanical forces play a complex and subtle role [5, 18–26]. On the one hand, local [27–30] and tissue-scale [31–35] mechanical stresses associated with cellular-shape changes directly drive morphogenetic movements such as furrow formation. On the other hand, there is emerging evidence that mechanical transduction and feedback effects work together with the genetic patterning system to harmonize mechanical cell activities, resulting in robust progression of morphogenesis [1–4, 32, 36–40].

Our paper is focused on a detailed analysis of mechanical processes that are responsible for formation of the cephalic furrow (CF) in the *Drosophila melanogaster* embryo (Fig. 1). CF is a transient morphogenetic structure on the boundary between the trunk and head patterning systems [41]. It has been hypothesized that CF plays a buffering role, mechanically isolating the head and trunk regions, and that it maintains tissue tension that supports mechanical signaling between morphogenetic movements [30].

**FIG. 1.**
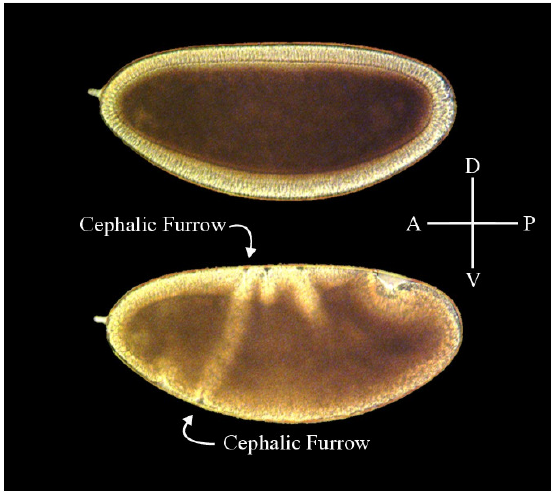
Cephalic furrow formation in the *Drosophila* embryo. Top image: sagittal plane before gastrulation, during the cellularization stage when embryo consists of a cellular monolayer. Bottom image: sagittal plane showing cellular movements and invaginations during gastrulation. The cephalic furrow is indicated (arrows). D–V: dorsal-ventral axis. A–P: anterior-posterior axis.

In contrast to the much better studied ventral furrow formation (VFF), the geometry and mechanics of CF formation (CFF) have been investigated only recently. Spencer *et al*. [42] provided the first detailed images of the furrow formation process and showed that initiator cell apices sink below the embryonic surface along a developing linear groove, followed by neighboring cells bending over to enter the groove, forming a cellular fold. The fold deepens as more cells enter, changing their shapes and pivoting as they move into the furrow [42]. More recently, our group [30] developed an advanced multinode lateral vertex model (MNLVM) to investigate the mechanics of CFF. The above study focused on the progression stage of CFF, during which subsequent cell pairs enter the furrow in a quasi-periodic process. Our analysis showed that the cells rolling over the furrow cleft lower their cellular pressure (and thus become more compliant) and that the high-pressure invaginated cells form a stiff block that provides support for apical-membrane tension forces that pull new cells into the furrow.

Such a supporting block, however, does not exist during the initiation phase of CFF. Thus, our present paper elucidates a different mechanism that initiates the CFF process. We provide evidence that the inward force that pulls the initiator cells toward the yolk sac is generated by tensile stress acting along the gradually expanding CF cleft on the curved embryo surface. We also argue that this tension is a source of mechanical signals that coordinate coherent furrow expansion and ensure its robustness.

We note that the existence of anisotropic tension along the CF was recently experimentally demonstrated by Eritano *et al*. [31], who showed that this tension helps align the initiator cells along the future CF cleft. It was also shown that tension acting along a narrow strip of a blastoderm can produce an invagination in a cell layer [34], and that it contributes to the generation of the ventral furrow (VF) [34, 35, 43]. The present study builds on these findings and on the results of our previous investigations of mechanical-feedback coordination of cell activities [26, 32, 44] to quantify how local cellular forces generated by apical, lateral, and basal cortices work together with long-range forces produced by actomyosin activity to ensure a coherent and robust CF invagination.

## II. THE GEOMETRY OF CEPHALIC FURROW AND KEY FORCES GOVERNING ITS FORMATION

The most important features of the initiation phase of CFF are presented in Figs. 2–4. Figures 2(a) and 3 show the *en face* view of a lateral side of the developing embryo, and Figs. 2(b) and 4 present the cross-sectional view of CF [also see the schematic in Figs. 2(c)]. In the following subsections, we present some crucial observations regarding the geometry and dynamics of CF initiation to set a groundwork for our analysis of mechanical processes that govern CFF.

**FIG. 2.**
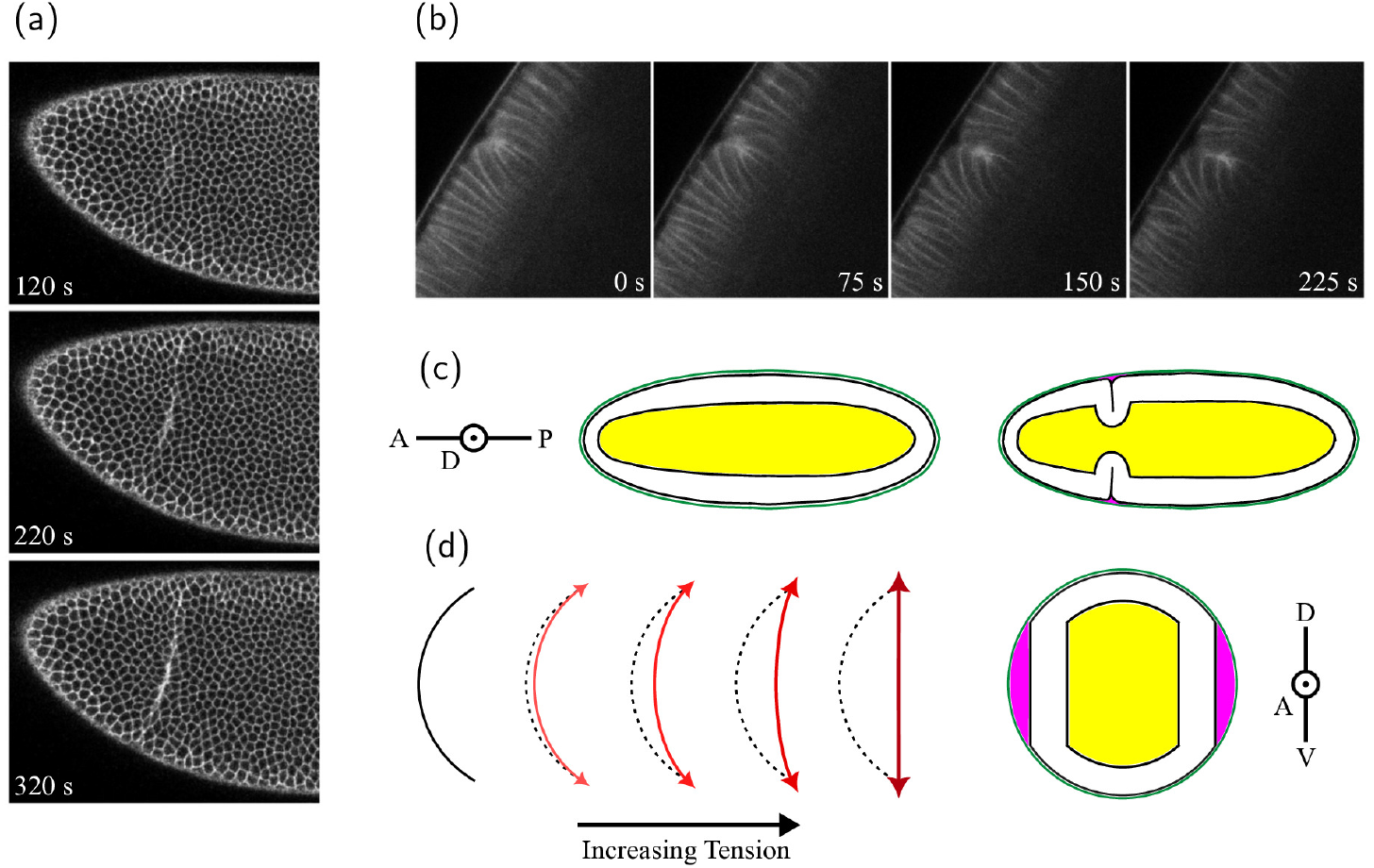
CF initiation and development. (a) *En face* and (b) cross-sectional confocal time-lapse of a Spider-GFP embryo show the initial emergence and subsequent expansion of the CF cleft. The vitelline membrane is visible in (b) as a thin white line above the apical surfaces of the cells. (c) Schematics before and during CFF. (d) The expansion of the CF cleft produces tension along the apical surface of the cell layer. Tension along the outer surface of a curved deformable medium will produce an effective tension-induced inward force, flattening the surface.

### A. *En face* view analysis

CF begins in two small lateral regions on the left and right sides of the embryo, and then gradually expands in the dorsal and ventral directions [Fig. 2(a)]. The expansion of the furrow involves transient heterogeneities manifested in Fig. 2(a) by an uneven fluorescent signal from membranes of activated initiator cells whose apices start to move away from the vitelline membrane and pass through the imaging plane (which is slightly below the embryo surface). The apices appear to taper as the initiator cells sink below the apices of neighboring cells.

A more detailed examination of the cell-shape pattern during the furrow expansion confirms that the activation of the initiator cells is spatially and temporarily heterogeneous [Fig. 3(a)]. The already activated cells form an expanding discontinuous chain with one-cell or two-cell gaps. The cells in the gaps activate after some delay, leading to formation of a continuous chain, which is the precursor of the furrow cleft. Similar heterogeneities in activation of initiator cells were reported by Eritano *et al*. [31].

**FIG. 3.**
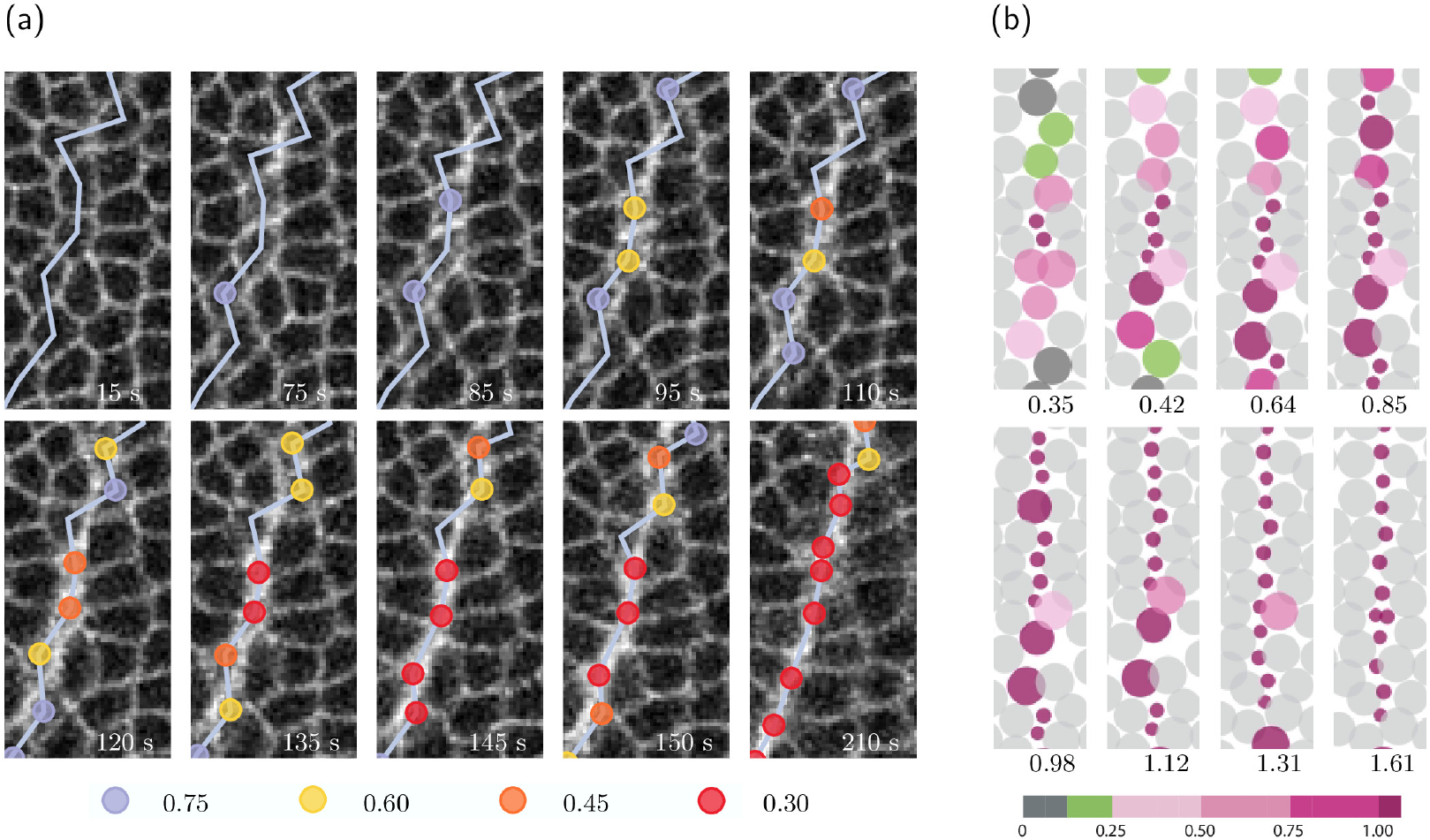
Development of a chain of activated initiator cells at the initial stage of CFF. (a) Processed *en face* confocal images of the lateral region where CF initiation occurs. The activated cells are marked by circles, color coded by the fractional reduction of the visible initiator-cell-apex width relative to its initial width (as labeled). The blue line indicates the final connectivity of the chain of activated initiator cells. (b) Formation of a chain of activated initiator cells, modeled as a stress-coordinated constriction process. The activated (constricted) initiator cells are represented by the small circles. The susceptible cells are color coded by the value of the triggering stress (8) normalized by the cutoff value *σ*_c_ (see Sec. III A for pertinent details of the model). The passive particles are depicted in light gray. The simulation-frame labels show the time normalized by the time at which 50% of susceptible cells have activated. The experimental images and the simulation frames present a blowup of the lateral region where the CF starts.

The spatiotemporal structure of the activation-chain patterns is closely similar to cellular-constriction chain (CCC) patterns that develop during the apical-constriction phase of VFF [32]. Since the CCC patterns originate from tensile-stress feedback correlating apical constrictions [32, 35, 44], we argue (see Sec. IV) that the inhomogeneities in CF initiation are a signature of tensile-stress feedback that guides CF expansion. This conjecture is consistent with the fact that tensile stress develops along the CF [31].

### B. Cross-sectional view analysis

The cross-sectional view of the initiation phase of CFF is presented in Figs. 2(b) and 4. The time-lapse sequence of confocal images of a live embryo [Fig. 2(b)] and the time-lapse montage of fixed embryo images (Fig. 4) show that CF invagination passes through several phases defined by morphological changes [42].

**FIG. 4.**
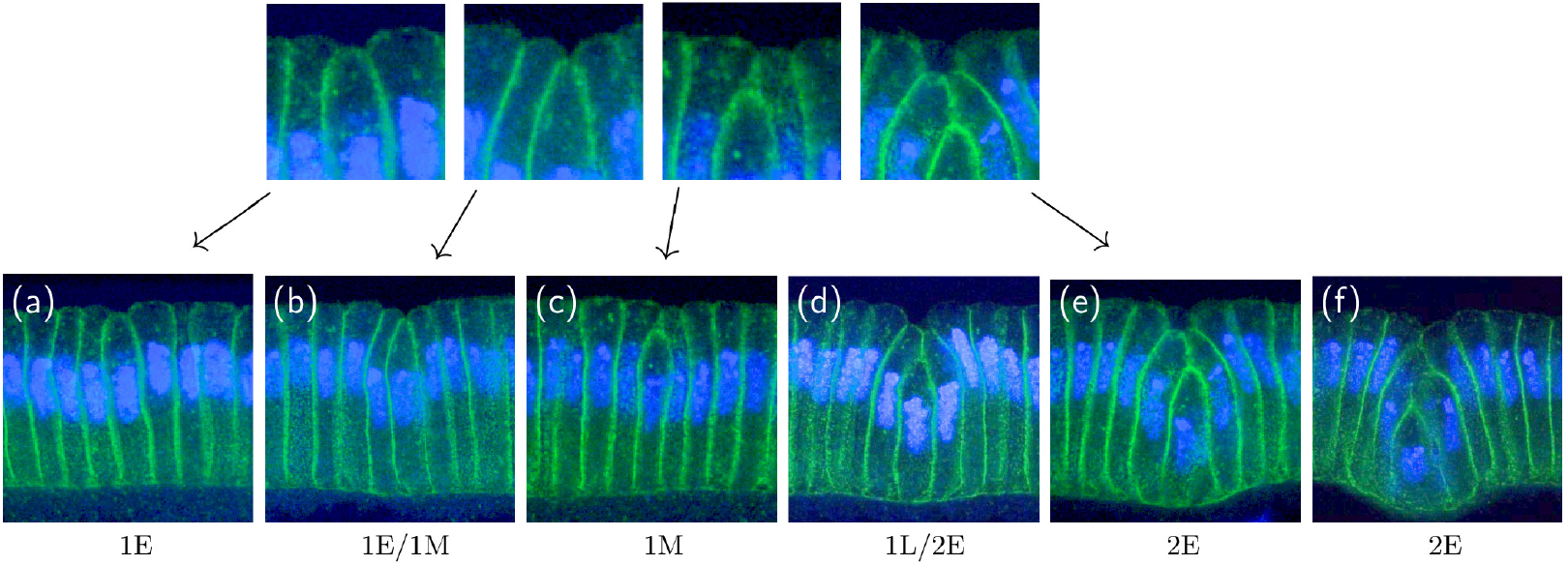
A time-lapse montage of contrast-enhanced cross-sectional fixed embryo images showing several stages of CF initiation. Membranes (Nrt, green) and nuclei (Hoechst, blue). (a) Phase 1E: the apical membrane of the initiator cell drops below the blastoderm apices. (b) Phase 1E–Phase 1M transition: the initiator cell shortens and adjacent cells bend into the furrow just before the apical membranes of adjacent cells meet. (c) Phase 1M: the apices of the adjacent cells are in contact and the initiator and adjacent cells enter the furrow. (d) Phase 1L–Phase 2E transition: the initiator cell has widened and shortened, and the top of the initiator cell nucleus has passed below the bottoms of the non-invaginated cell nuclei. The initiator cell and its neighboring cells’ bases start to protrude into the yolk sac. (e,f) Phase 2E: subsequent cell pairs descend into the furrow. The initiator cell continues cell shortening and basal expansion, as do the subsequent cells. The CF protruded deeper into the yolk sac. Magnified views of the CF cleft region are shown in the insets.

In Phase 1E (first phase–early) the apices of the initiator cells drop below the neighboring blastoderm apices and the initiator cells shorten. A small perivitelline-fluid region forms at the emerging CF cleft. Phase 1M (first phase–middle) begins when the apices of the adjacent cells contact each other over the initiator cell apices. Phase 1M is a newly defined part of Phase 1 that is associated with a tension-induced inward force. Phase 1L (first phase–late) starts when the tops of the initiator cell nuclei pass below the bottoms of the blastoderm nuclei. Phase 2E (second phase–early) starts when the bases of the initiator cells protrude below the baseline of the blastoderm into the yolk sac. The transition to phase 2L (second phase–late) takes place when the apices of the initiator cells drop below the bases of the other cells of the blastoderm [42].

The descent of the apical end of the initiator cell into the newly forming groove requires an inward-directed force to balance the yolk pressure, which pushes the epithelial layer against the vitelline membrane. As shown in Sec. V, the tension of lateral membranes cannot produce such a force without causing a deformation of the basal surface of the epithelial layer toward the vitelline membrane. A concave shape of the epithelial layer, however, is not observed in the *Drosophila* embryo. Thus, it follows that an additional, previously unrecognized, inward-directed force must be contributing to the initiation of the CF invagination.

This force is provided by the tension *τ* acting along the furrow [Fig. 2(d)]. Existence of tension *τ* in a narrow strip along the emerging furrow cleft was demonstrated by Eritano *et al*. [31]; as discussed in Sec. II A, this tension is involved in coordinating activation of initiator cells. Since tension *τ* operates over the curved surface of the embryo, it exerts the inward force

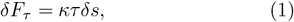

which pushes the epithelial layer into the CF cleft. Here *κ* is the surface curvature along the CF and *δs* is the arclength element on which the force *δF*_*τ*_ acts. So far, an analogous inward force has been demonstrated for VFF [34, 35, 43] but not for CFF.

Our results strongly point to the conclusion that the tension-generated inward force (1) is likely a universal mechanism involved in furrow generation. In what follows we argue that the tension *τ* and the associated force *δF*_*τ*_ not only mechanically shape the epithelial layer, but also produce mechanical signals that coordinate cell activities on the local and embryo-scale levels.

## III. NUMERICAL MODELS OF CFF MECHANICS

To shed light on the role of mechanical forces, mechanical feedback, and geometrical cell-shape transformations in CF initiation, we have developed two complementary numerical models [Fig. 5(a)]. These models analyze two different aspects of the CFF mechanics: the forces propagating along the furrow (the *en face* force-center model, FCM), and the forces that are orthogonal to the furrow direction (the multi-node lateral vertex model, MN-LVM).

**FIG. 5.**
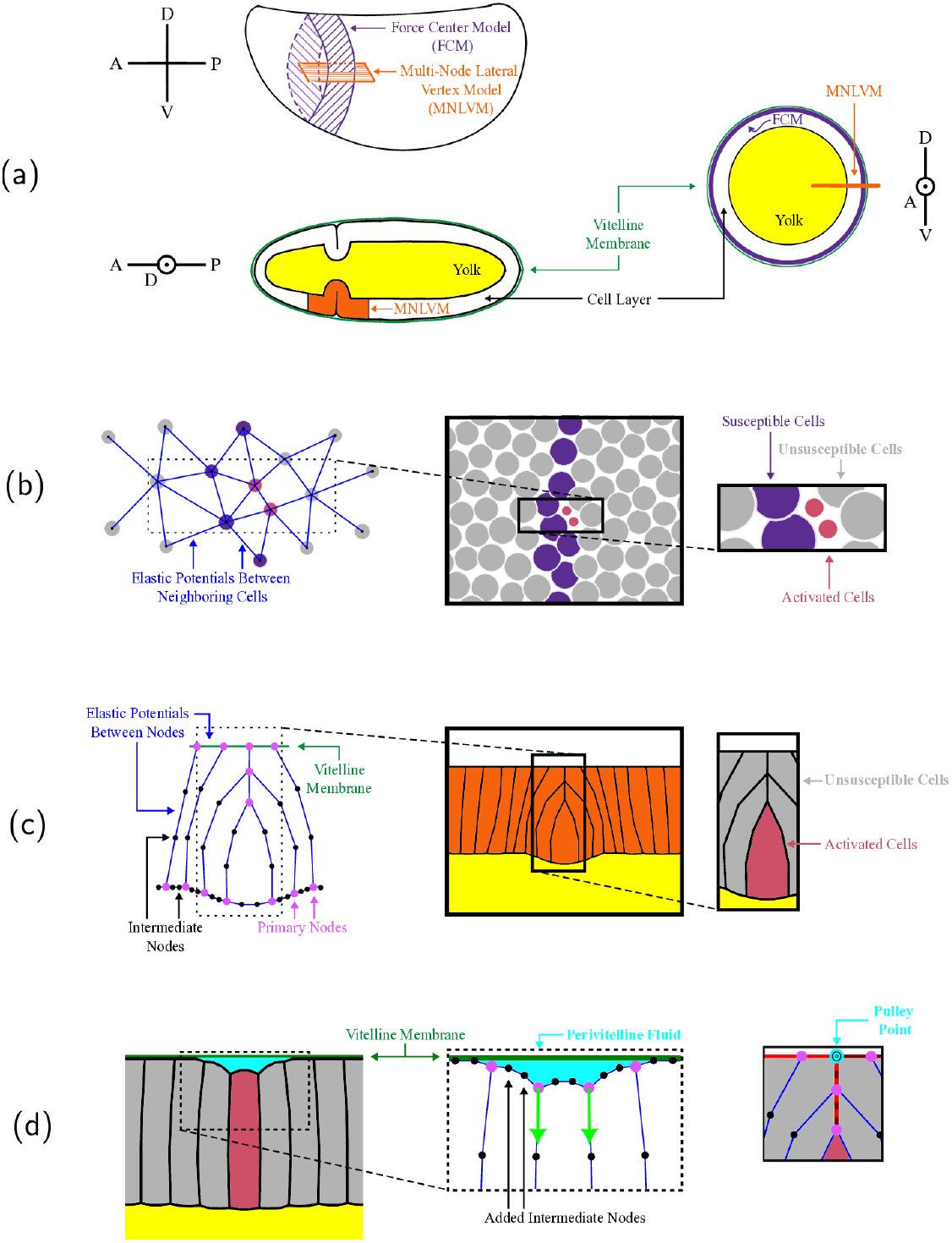
Main features of the *en face* force-center model, FCM, and multi-node lateral vertex model, MNLVM. (a) Schematics highlighting embryo areas considered by each model. FCM focuses on the apical surface of the region along which the CF cleft expands. MNLVM concentrates on a cross-section of the cell layer which includes the initiator cell(s) and other cells that will roll over the furrow cleft as the CF deepens. (b) FCM describes how initiator cells are activated as the CF cleft expands. Cells are treated as force centers that interact with their immediate neighbors via elastic potentials (blue lines). Susceptible cells (purple) are capable of becoming activated (puce) while unsusceptible cells (gray) are not. (c) MNLVM represents cells as a collection of vertices that interact through elastic potentials (blue lines). Adding intermediate vertices (black) between the primary vertices (pink) introduces a new degree of freedom by allowing membranes to bend as a result of pressure variations across them. (d) During the initiation phase, the perivitelline fluid plays a vital role and must be explicitly represented to capture realistic cellular deformations. Intermediate apical vertices are introduced along with a downward force on the primary vertices of the initiator cell to mimic the tension-induced inward force. To describe the subsequent evolution, we use an implicit representation of the perivitelline space by introducing a pulley point which guides the apical membranes of cells that meet as they roll over the furrow cleft.

Our first model, the *en face* FCM [Fig. 5(b)], focuses on the interrelation between the tensile stress propagating along the furrow and the activation of initiator cells via mechanical feedback. This model is based on our active-granular-fluid (AGF) approach that was used to describe apical constrictions coordinated via tensile stress during the early slow phase of VFF [32]. Based on results shown in Fig. 3, we argue in Sec. IV that similar feedback and stress propagation mechanisms are active during CFF.

Our second approach, MNLVM, describes the cross-sectional view of the furrow [Fig. 5(c)], and it incorporates membrane curvatures into the analysis of the furrow mechanics. [30], The membrane curvatures provide vital information regarding pressure differences from cell to cell and yield additional degrees of freedom that are crucial for assessing whether the model correctly describes mechanical processes governing the furrow dynamics.

The MNLVM is used here in two versions: our original technique [30], in which the perivitelline fluid region is modeled implicitly [Fig. 5(c)] and a new enhanced method with an explicit representation of the perivitelline fluid [Fig. 5(d)].

### A. Force-center model of furrow expansion

According to recent investigations of CF development, there is an accumulation of myo-2 at the adherens junctions near the apical surface of initiator cells at the onset of activation ([31, 42]). F-actin accumulates along the apical and subapical regions of the initiator cells ([42]). The associated actomyosin activity generates tensile stress propagating along the furrow [31]. To model the heterogeneous tensile-force propagation and mechanical feedback generated by the stress, we use a coarse-grained FCM [Fig. 5(b)], in which each individual force center represents an apical end of a cell and the underlying actomyosin machinery.

The force centers *i* and *j* that represent adjacent cells interact via a spring potential

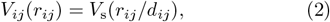

where *r*_*ij*_ is the distance between centers *i* and *j*, 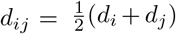 is the average potential range of the force centers *i* and *j*, and

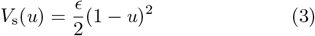

is a harmonic potential that depends o the normalized interparticle distance *u* = *r*_*ij*_*/d*_*ij*_.

The parameter *ϵ* in Eq. (3) sets the energy scale for the interparticle interaction. The potential (2) includes both the repulsive and attractive parts, associated with cell elasticity and adhesion. The cells that in the initial state are not adjacent interact only via the repulsive part of the spring potential,

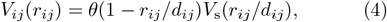

where *θ*(*x*) is the Heaviside step function. The interaction force between force centers *i* and *j* is

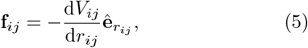

where 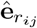 is the unit vector in the radial direction. The potential range *d*_*i*_ corresponds to the diameter of the cell *i* in the absence of forces exerted by the surrounding cells.

The generation of contractile forces by the actomyosin activity is represented by the reduction of the potential range of the activated force centers

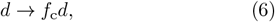

where *f*_c_ *<* 1 is the constriction factor. (In our simulations we use *f*_c_ = 0.4). The constrictions (6) occur only in a narrow strip of susceptible cells [Fig. 5(b)], corresponding to genetically predefined zone of CF activation. Based on *in vivo* measurements by Eritano *et al*. [31], we choose the width of this zone, normalized by the average particle diameter, to be Δ = 1.4.

Due to the presence of adhesion forces [the attractive part of the potential (3)], the effective diameter reduction (6) generates tensile stress that affects surrounding cells. Since the constrictions occur in the narrow band, the tensile stress propagates along the emerging furrow cleft. The associated mechanical feedback is incorporated in our model by making the particle-constriction probability per simulation step tensile-stress sensitive.

Following our previous analyses of the apical-constriction process in VFF [32, 35], the stress feedback is described via the feedback parameter

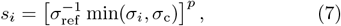

where *σ*_*i*_ is the triggering tensile stress exerted on cell *i* by the adjacent cells, parameter *p* defines the stress-response profile, *σ*_c_ is the cutoff value above which the stress feedback saturates, and *σ*_ref_ is the normalization factor. In our simulations we use *p* = 3, *σ*_c_ = 0.7*σ*_ref_, and *σ*_ref_ is defined as the average tensile stress experienced by a single cell constricted in the initial configuration.

Since the actomyosin network that forms during CF initiation is limited to a narrow band oriented along the emerging furrow, we assume that the triggering stress *σ*_*i*_ includes only the tensile part of the longitudinal stress-tensor components

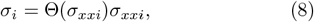

where

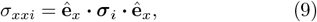

**ê**_*x*_ is the unit vector oriented along the furrow, and

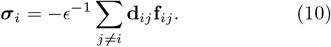

Here 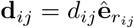, where 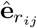 is the unit vector oriented from particle *j* to particle *i*.

The constriction probabilities per simulation step *P*_*i*_(*s*_*i*_) are calculated from the relation

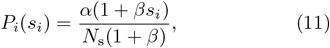

where the parameter *β* measures the magnitude of the stress-feedback contribution relative to the stress-insensitive background value. The normalization factor *α/N*_s_ (where *N*_s_ is the total number of susceptible particles, including the unconstricted and constricted ones) controls the average number of constrictions in each step of the stochastic simulation process.

In each simulation step, the system is first equilibrated, the triggering stress is evaluated from Eq. (8), and the force-center diameters *d*_*i*_ are reduced with the probability (11). Subsequently, the system is equilibrated again. In our simulations we use *α* = 1.0 and *β* = 5.0 *×* 10^4^. The large value of *β* ensures that random uncorrelated constrictions are much less likely than stress-correlated ones. For simplicity, we apply a one-step diameter reduction process. A gradual initiator-cell activation can be implemented using the approach described in our previous study [35],

Our simulations are performed in a rectangular domain 80-particle long and 24-particle wide. In a system with the susceptible-particle-zone of width Δ = 1.4 there are approximately *N*_s_ = 100 particles. We apply the periodic boundary conditions in the furrow direction *x*, consistent with the fact that CF develops into a closed loop along the entire embryo perimeter. In the transverse direction we use boundary conditions that mimic the elasticity of the surrounding epithelial tissue [35].

### B. Multi-node lateral vertex model

#### 1. Multi-node membrane representation

To describe cross-sectional dynamics of CF initiation, we use MNLVM that was successfully applied to analyze the later progression stage of CFF [30]. In MNLVM, each cell is described by a set of primary and intermediate nodes. The primary vertices define the meeting points of the apical, lateral and basal membranes. The intermediate nodes divide each membrane into equal-size segments to reflect the fact that the membranes are deformed by cross-membrane pressure jumps [Fig. 5(c)].

In our presentation below, all relations are given in a dimensionless form. We choose the length scale *L*_0_ and the energy scale *E*_0_ as primary units. Accordingly, all lengths are scaled by *L*_0_, potential energies by *E*_0_, areas by 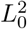, membrane tensions and forces acting on vertices by *σ*_0_ = *E*_0_*/L*_0_, and pressures by *σ*_0_*/L*_0_. This normalization is simpler than the one employed in our earlier study [30].

The system is described using Cartesian coordinates (*x, y*), with *x* parallel to and *y* normal to the epithelial layer. The CF cleft is at the position *x* = 0, the vitelline membrane is at *y* = 0, and *y <* 0 corresponds to the embryo interior. The position of node *i* is denoted by (*x*_*i*_, *y*_*i*_).

#### 2. Membrane tensions, cell pressures, and blastoderm tension

The membrane segments are described by linear-spring potentials

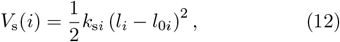

where *i* is the index of a segment, *l*_*i*_ and *l*_0*i*_ are its actual length and rest length, and *k*_s*i*_ is the spring constant. The energy associated with the cross-sectional area of the cells is

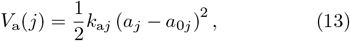

where *a*_*j*_ and *a*_0*j*_ are the actual and rest area of cell *j*, and *k*_a*j*_ is its area elasticity constant. The membrane tensions and cell pressures associated with the potentials (12) and (13) are

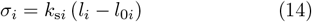

and

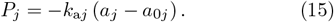

The effect of the epithelial tissue outside the explicitly simulated region on the invagination mechanics is incorporated via forces acting on the primary boundary nodes of the explicitly simulated domain. The potential of the boundary force acting on the boundary node *n* is

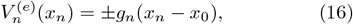

where *g*_*n*_ is the strength of the force acting on node *n, x*_0_ is the reference position, and the plus (minus) sign applies to the cells on the left (right) boundary of the simulation domain.

The boundary force mimics the elastic interaction between the explicitly and implicitly simulated parts of the epithelial tissue. The total force

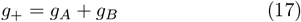

corresponds to blastoderm tension (*g*_+_ *>* 0) or compression (*g*_+_ *<* 0), and

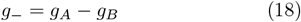

is associated with the boundary shearing stress, which controls the tilt of lateral membranes. In the above expressions the subscript *A* refers to the apical and *B* to the basal boundary nodes.

#### 3. Implicit and explicit representations of the perivitelline fluid region

In our original version of the MNLVM the apical membranes of non-invaginated cells are constrained to move along the line representing vitelline membrane. The perivitelline fluid region is modeled in a simplified way, as a pulley point over which apical membranes of invaginating cells roll over when they enter the furrow [30]. In our present study this implicit representation of the perivitelline fluid is used in modeling of the later part of the CF initiation process (phases 1L and 2E in the nomenclature explained in Sec. II B [42]). However, to determine mechanical forces that govern initial descent of the initiator cells into the furrow (phase 1E), the perivitelline fluid region needs to be explicitly modeled.

In this explicit approach the positions of the primary and intermediate apical nodes are not constrained to the line representing the vitelline membrane. The node– vitelline membrane overlap is prevented by a short-range repulsive potential of a generalized Yukawa form,

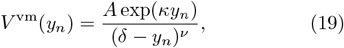

where *A, κ, δ*, and *ν* are numeric parameters.

The effect of the negative pressure of perivitelline fluid, which hinders the separation of the apical nodes from the vitelline membrane, is accounted for by the perivitelline-fluid area potential

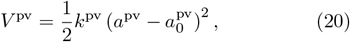

where *a*^pv^ and 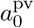 denote the actual area and rest area of the region between the apical cell membranes and vitelline membrane, and *k*^pv^ is the corresponding elasticity constant.

In our simulations, the unperturbed cells at the boundary of the simulation domain are rectangular and have the aspect ratio *r* = 5, where *r* = *L*_ab_*/W*, and *W* and *L*_ab_ denote cell width and the apicobasal cell length, respectively. To accommodate membrane curvatures, membranes are divided into three equal-size segments.

In the normalization where the width of an unperturbed cell is chosen as the characteristic length *L*_0_ (i.e., for the dimensionless width *W* = 1) we use the following parameter values: *k*_s*i*_ = 4.5 for the spring constants of the apical and basal segments, *k*_s*i*_ = 0.9 for spring constants of the lateral segments, and *k*_a*j*_ = 0.2 for the area spring constants. In simulations with explicit perivitelline-fluid region we also use 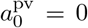 and *k*^pv^ = 0.2 in Eq. (20) and 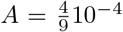, *κ* = 45, *δ* = 0.2, and *ν* = 2 in Eq. (19).

To accelerate numerical convergence of our optimization procedures toward experimentally determined geometry under investigation, in some simulations with pulley-point representation of perivitelline-fluid region we use modified values of *k*_s*i*_ and *k*_a*j*_. As explained in Ref. [30], this change does not affect the final optimal structures, because the same state can be obtained using different combinations of spring constants and rest lengths and areas.

#### 4. Tensile-stress-generated inward force

According to experimental evidence [31], the tensile stress producing the inward force (1) is concentrated near adherens junctions in the lateral membranes near the apical cell surface. Since in our model the junctions are not explicitly modeled, the tensile-stress-generated compressing force *δF*_*τ*_ is applied to the primary apical vertices of the initiator cells [Fig. 5(d)]. The force on node *n* is included by adding the corresponding potential

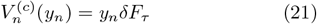

to the potential of the vertex model, where *y*_*n*_ is the coordinate normal to the vitelline membrane, pointing outward.

#### 5. Equilibrium configurations of the vertex model

In our comparison of the model with experimental images it is assumed that the system evolves through a set of equilibrium configurations determined by the force balance at each node. The forces are generated by membrane tensions and cell pressures, boundary forces, and the force *δF*_*τ*_. Since the viscosity of cytoplasm is of the order of 1 Pa s [45], the stresses associated with membrane tensions and cell pressures are of the order of 10 Pa, and the shear rate in the invagination region is of the order of 10^−2^ s^−1^ [30], the quasistatic-evolution assumption is justified.

As discussed in our earlier study [30], the MNLVM has more degrees of freedom than the number of model parameters that we can control. Therefore, only a subset of geometrically allowable vertex configurations is achievable for a system in mechanical equilibrium. Since there is no guarantee of agreement between the experimentally observed and simulated configurations, the model allows us to test whether all key forces that govern the system mechanics are correctly accounted for (i.e., a lack of such agreement would imply that some important mechanical contributions are missing). We can also evaluate the relative importance of different mechanical contributions at different stages of furrow formation.

## IV. TENSILE-STRESS COORDINATION OF INITIATOR-CELL ACTIVATION EVALUATED USING FCM APPROACH

### A. Formation of initiation chains

In this section we discuss simulation results obtained using our *en face* FCM of mechanical-feedback coordination of initiator-cell activation. The numerical results are discussed in the context of experimental findings by our group and others [30, 31, 46].

The simulated process starts by constricting several seed particles in two regions corresponding to the small lateral domains where the CF initiation begins. The subsequent initiation events, modeled as particle constrictions (which represent actomyosin contractions), are tensile-stress coordinated (as described in Sec. III A).

A time-lapse sequence of simulation frames showing the expansion of the CF initiation domain is presented in Fig. 6. The activated initiator cells are represented by small circles, with the diameter commensurate with the reduced interaction-potential range [Eqs. (2)– and (6)]. The particle color shows the magnitude of the triggering tensile stress, with deep red indicating the strongest tension.

**FIG. 6.**
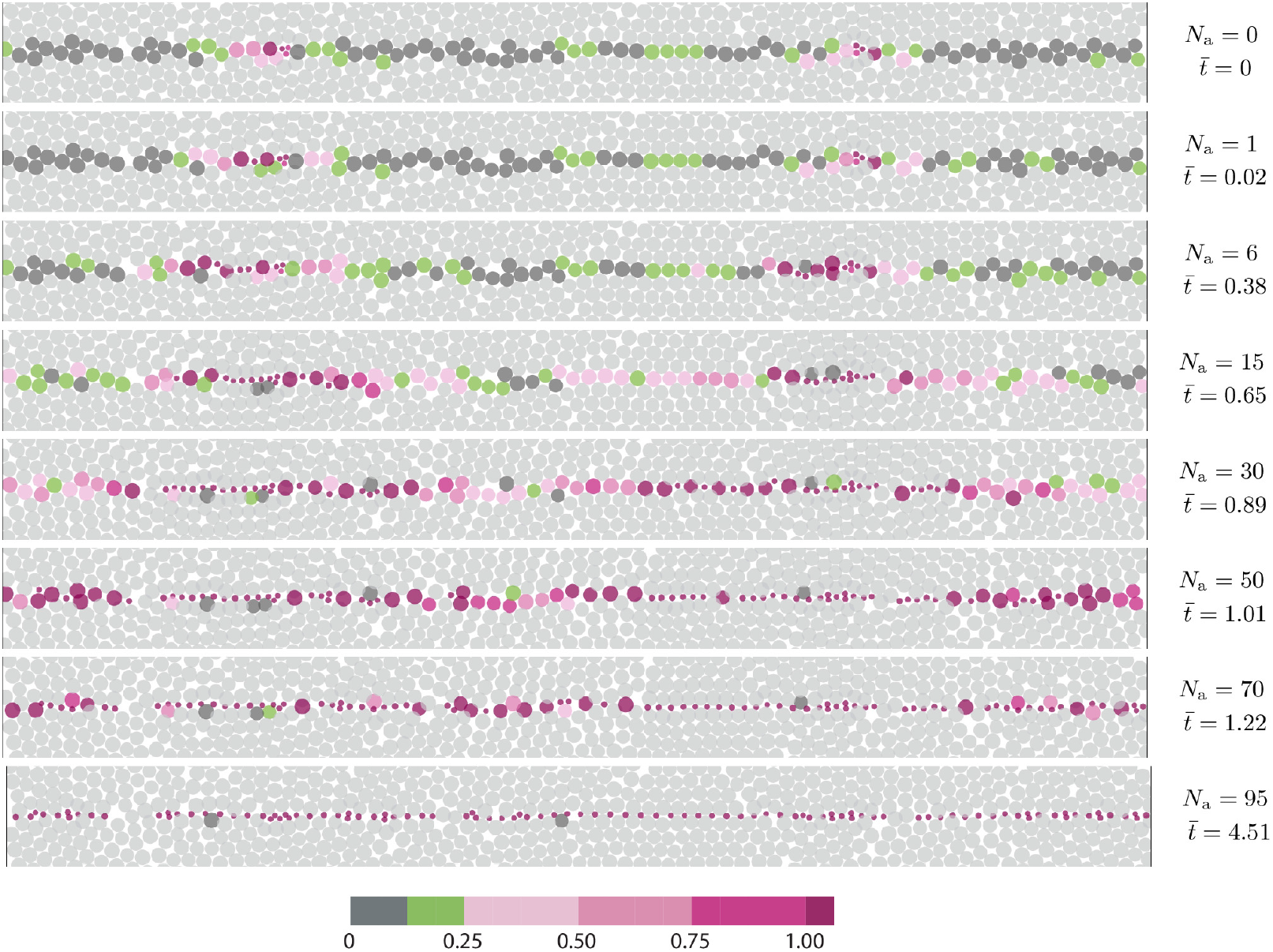
The initiator-cell activation wave coordinated by tensile stress along the developing CF cleft. In the initial simulation frame 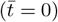 two three-particle clusters of constricted seed cells (marked by small red circles) are present in two lateral regions where the initiation process starts. The passive cells are represented by gray circles and initiator cells by colored circles, with the color representing the value of the triggering stress (8) normalized by the cutoff value *σ*_c_. The time-lapse frames are labeled by the number of constricted non-seed cells *N*_a_ and by the time 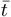 normalized by the time at which 50% of susceptible non-seed particles have activated The subsequent frames show the buildup of triggering tensile stress and heterogeneous expansion of the initiator-cell activation chains.

The consecutive panels in Fig. 6 are labeled by the number of constricted (activated) non-seed particles *N*_a_. The rescaled time 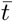 (proportional to the number of simulation cycles) is normalized by the time at which 50% of susceptible non-seed particles have activated. The time 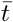 is set to zero when only the seed particles have constricted (i.e., for *N*_a_ = 0).

The results in the top panel (*N*_a_ = 0) show that constriction of the seed particles generates tensile stress in their neighborhood; the stress propagates over a distance of several particle diameters along the furrow. This tensile stress triggers constrictions of subsequent particles, leading to further enhancement of the tension. The constrictions cause a gradual expansion of the initiated region along the precursor stress line, until most of the susceptible particles have constricted.

Due to the non-local character of stress propagation, the stress enhancement of the constriction probability results in the spatiotemporal heterogeneity of the constriction process, reflected in discontinuous constriction chains. A blowup of the region near the point where the furrow expansion begins [Fig. 3(b)] shows that the simulated constriction pattern is qualitatively similar to the one observed *in vivo* [Fig. 3(a)]. The details of the pattern vary from simulation to simulation [Figs. 3(b) and 6 show results for different initial conditions but the same simulation-cell size], but the discontinuous character of the expansion process remains qualitatively the same.

The constriction/initiation heterogeneities occur not only at short times but they persist through the entire initiation process. A similar heterogeneous initiation pattern was reported for the *Drosophila* embryo by Eritano *et al*. [31]. Moreover, the simulated fraction *f*_a_ = *N*_a_*/N*_s_ of the susceptible cells (*N*_s_) that have already activated (*N*_a_) (Fig. 7) has a similar time dependence to the activation profile reported for *Drosophila* [31]. Both in simulations (Fig. 7) and *in vivo* (Fig. 1(c)] of [31]) the activation rate is approximately constant for 0.2 ≲ *f*_a_ ≲ 0.8, decreases for *f*_a_ ≳ 0.8, undergoes large temporal fluctuations, and there is a random variation between different embryos.

**FIG. 7.**
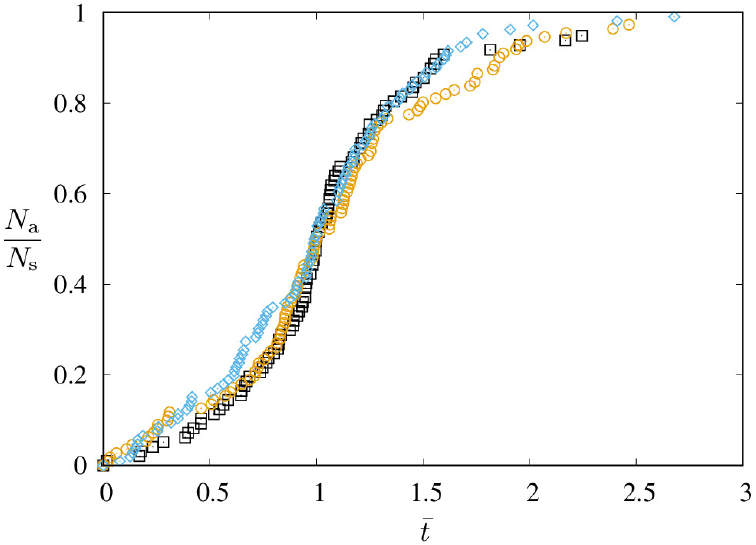
FCM predictions for the fraction of activated initiator cells vs the normalized time 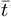 in a system with tensile-stress correlated initiation events. Different symbols correspond to three different initial conditions. The predicted time profile of the initiation events is similar to the corresponding *in vivo* activation-fraction result reported for the *Drosophila* embryo in Fig. 1(c) of [31].

### B. Initiation robustness

The agreement between the simulated and experimental initiation patterns furnishes a strong argument that tensile stress provides the essential signal that guides the furrow expansion. We will now discuss how mechanical feedback enhances the robustness of the initiation process.

To this end, we compare the stress-guided CF initiation (Fig. 6) with constrictions guided by the presence of initiated neighbors (Fig. 8). In both cases the process starts from the same initial configuration. In the latter (neighbor-triggering) case, we use the feedback parameter

**FIG. 8.**
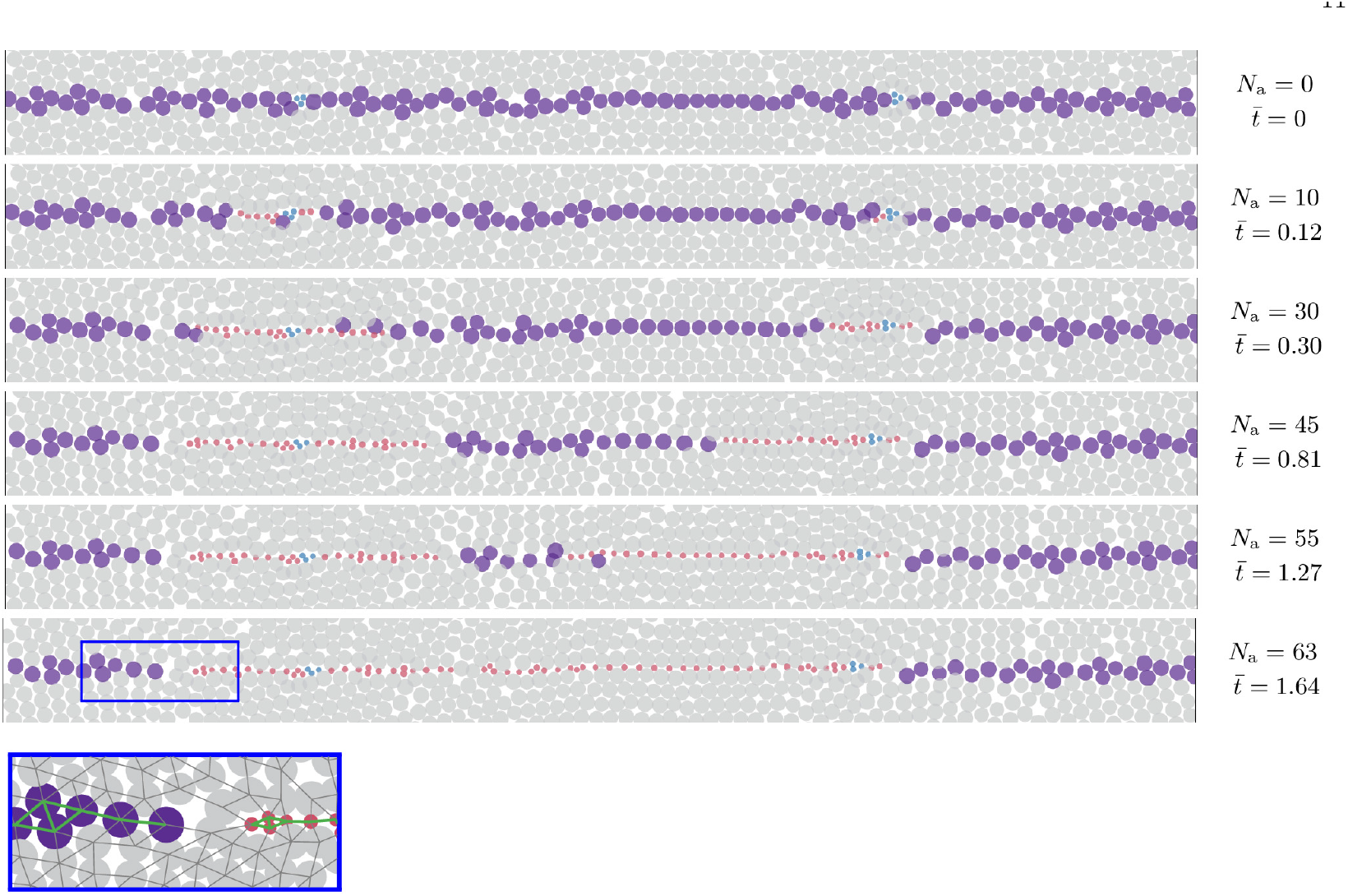
The initiator-cell activation wave triggered by near-neighbor interactions. The initiator cells activate if they have at least one already activated connected neighbor. The non-activated and activated initiator cells are represented by large purple and small red circles, respectively, and non-initiator cells are marked by gray circles. The initial condition at 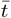 is the same as the one used in Fig. 6. The initially activated seed particles are indicated in blue. The time-lapse simulation frames show that the wave propagation can be arrested at stochastically occurring topological defects, where the network of connections between initiator cells has a discontinuity. The inset shows the topological defect in the region marked by the blue box in the last panel 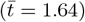. The neighbor connections between the initiator cells are marked in the inset in green, and the remaining connections in dark gray.

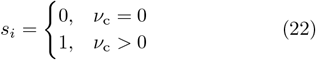

in Eq. (11), where *ν*_c_ is the number of the nearest neighbors of cell *i* that have already constricted.

Figure 8 shows that at first the neighbor-triggered initiation chain expands, without, however, forming transient gaps, observed both *in vivo* and in simulations of stress-guided initiation. The expansion continues until the front of the chain reaches a position where the graph of the nearest-neighbor connections between the susceptible cells has a discontinuity. At such a topological defect in the susceptible-cell connectivity network the expansion stops, and the process cannot proceed without employing alternative modes of intercellular communication. In contrast, the stress-correlated chain expansion continues (Fig. 6), owing to the buildup of tensile stress on the other side of the defect.

In a recent study Popkova *et al*. [46] showed that the expansion of CF can be viewed as a mechanical trigger wave propagating along a genetic guide and hypothesized that the wave is generated by stresses associated with apicobasal shortening of neighboring cells. However, such stress, which is oriented primarily in the apicobasal direction, has a much shorter range than the tensile stress along the furrow, and therefore would not ensure a robust invagination. Our model also describes the lateral furrow expansion as mechanically based trigger wave that propagates along the genetic anterior-posterior pattern, but the mechanical guidance for the furrow expansion is provided by tensile stress, which can propagate across defects in the initiator cell network. Thus the tensile-stress guidance is more robust than guidance mechanisms based on short-range intercellular interactions.

## V. CEPHALIC-FURROW INITIATION ANALYZED USING MNLVM APPROACH

Section IV focused on the expansion of the initiation zone along the emerging furrow. We now analyze mechanical forces that cause the initiator and subsequent cells to dive into the furrow.

In what follows, the membrane tensions and applied forces are normalized by the tension 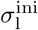 of the lateral membrane of the initiator cell in the initial state with no boundary and inward forces. In particular, we use the following notation

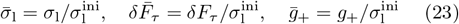

for the normalized lateral tension of the initiator cell 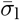, the tension-induced inward force 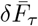, and the tension of the epithelial layer on both sides of the furrow 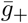. We also consider the relative reduction of the length of the lateral membrane of the initiator cell,

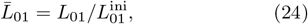

where *L*_01_ is the rest length of the initiator-cell lateral membranes in the current state and 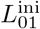 is their length in the unperturbed initial state.

### A. An analysis of the onset of the CF initiation process

#### 1. Formation of perivitelline fluid region at the CF cleft

Figure 9(a) shows an image of a fixed embryo during phase 1E of CF initiation. The apical membrane of the activated initiator cell has begun to move away from the vitelline membrane, creating a perivitelline fluid region (indicated by the arrow) above the apex of the initiator cell.

**FIG. 9.**
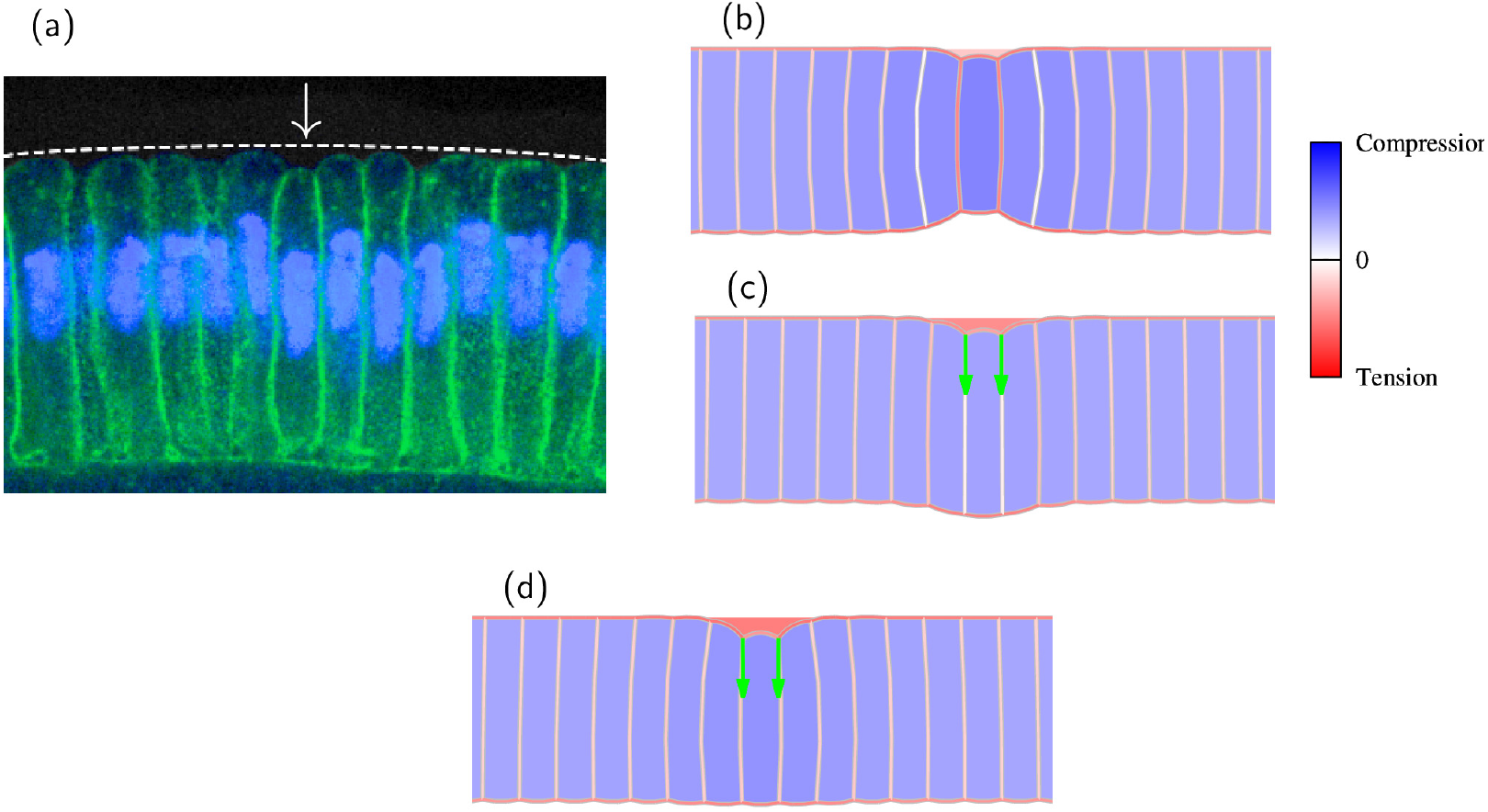
The role of tension-induced inward force 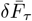 in CF initiation. (a) Contrast-enhanced confocal image of a fixed embryo showing formation of a perivitelline fluid region between the vitelline membrane and the apical membranes of the initiator and adjacent cells. Membranes (Nrt, green) and nuclei (Hoechst, blue). The approximate position of the vitelline membrane is marked with the dashed line and the perivitelline fluid region is indicated with an arrow. (b–d) MNLVM predictions for generation of the perivitelline fluid region. In (b) the perivitelline fluid region results from apicobasal shortening of the initiator cell by a factor 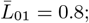 in (c) the perivitelline fluid region forms as a result of the tension-induced inward force 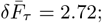 and the configuration shown in (d) is shaped by a combined inward force 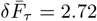 and apicobasal shortening with 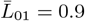. The inward force 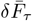 in (c) and (d) is indicated by the arrows.

The large curvature of the apical membranes of the initiator and adjacent cells and the moderate curvature of basal membranes indicate that the perivitelline fluid pressure is significantly lower than the cell pressure and that the cell and yolk pressures are approximately the same. Thus, the yolk pressure pushes the cell layer toward the vitelline membrane, and the active mechanical forces that generate CF must overcome the pressure difference between the yolk and perivitelline fluid.

Mechanical forces that drive the apical membrane of the initiator cell away from the vitelline membrane and cause formation of the perivitelline-fluid region are elucidated by results of our numerical simulations, shown in Figs. 9–11. These forces include the inward force 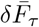 generated by the tension along the CF cleft, Eq. (1), and the tension 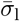 of the lateral membranes of the initiator cell. The simulations were performed using MNLVM with the explicit representation of the perivitelline fluid. Figures 9 and 10 present simulation results for a system with no blastoderm tension in the epithelial layer outside the invagination region,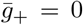. The influence of the blastoderm tension on CFF is discussed in Fig. 11.

**FIG. 10.**
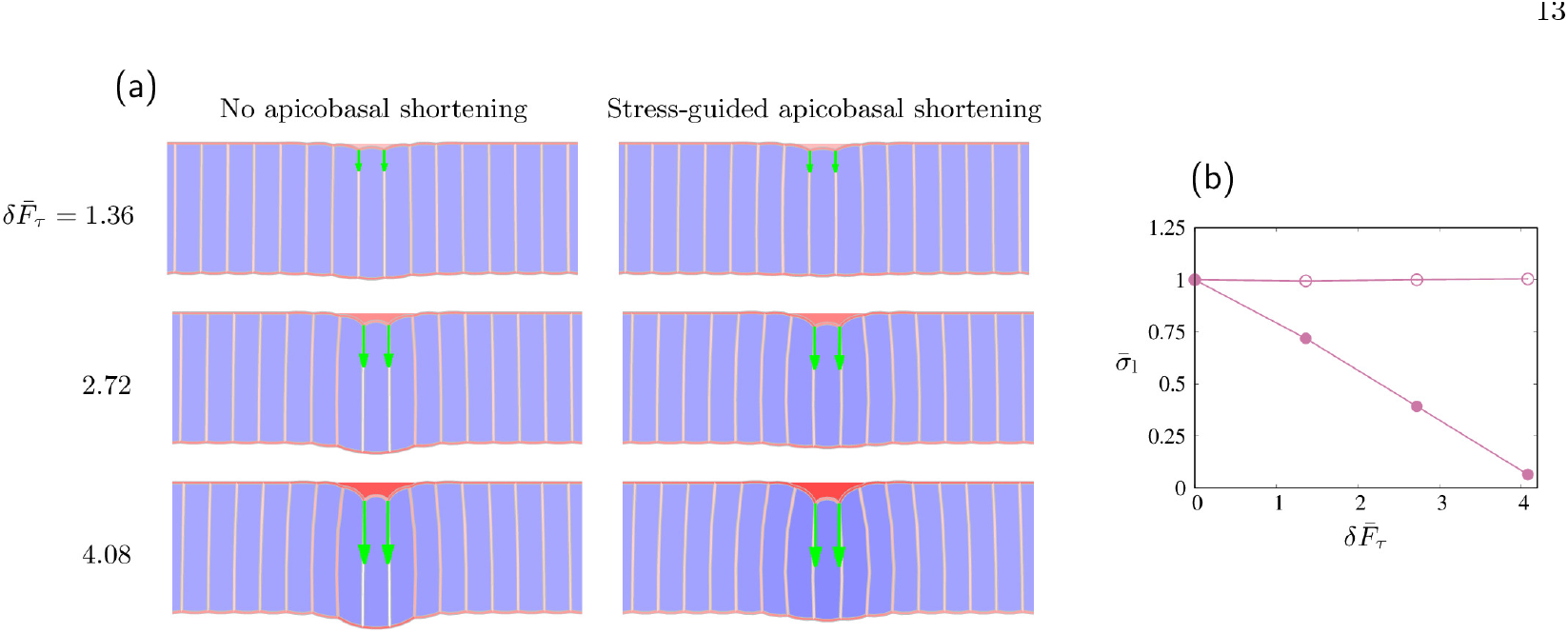
MNLVM analysis of inward-force-controlled initiator-cell shortening during phase 1E of CFF. (a) Cell configuration for different values of the normalized inward force 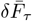 (as labeled) with no apicobasal shortening (left) and with stress-controlled apicobasal shortening that maintains the constant value of the lateral-membrane tension of the initiator cell (right). (b) Normalized lateral tension 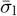 of the initiator cell vs the normalized inward force for a system with (closed circles) and without (open circles) apicobasal shortening.

**FIG. 11.**
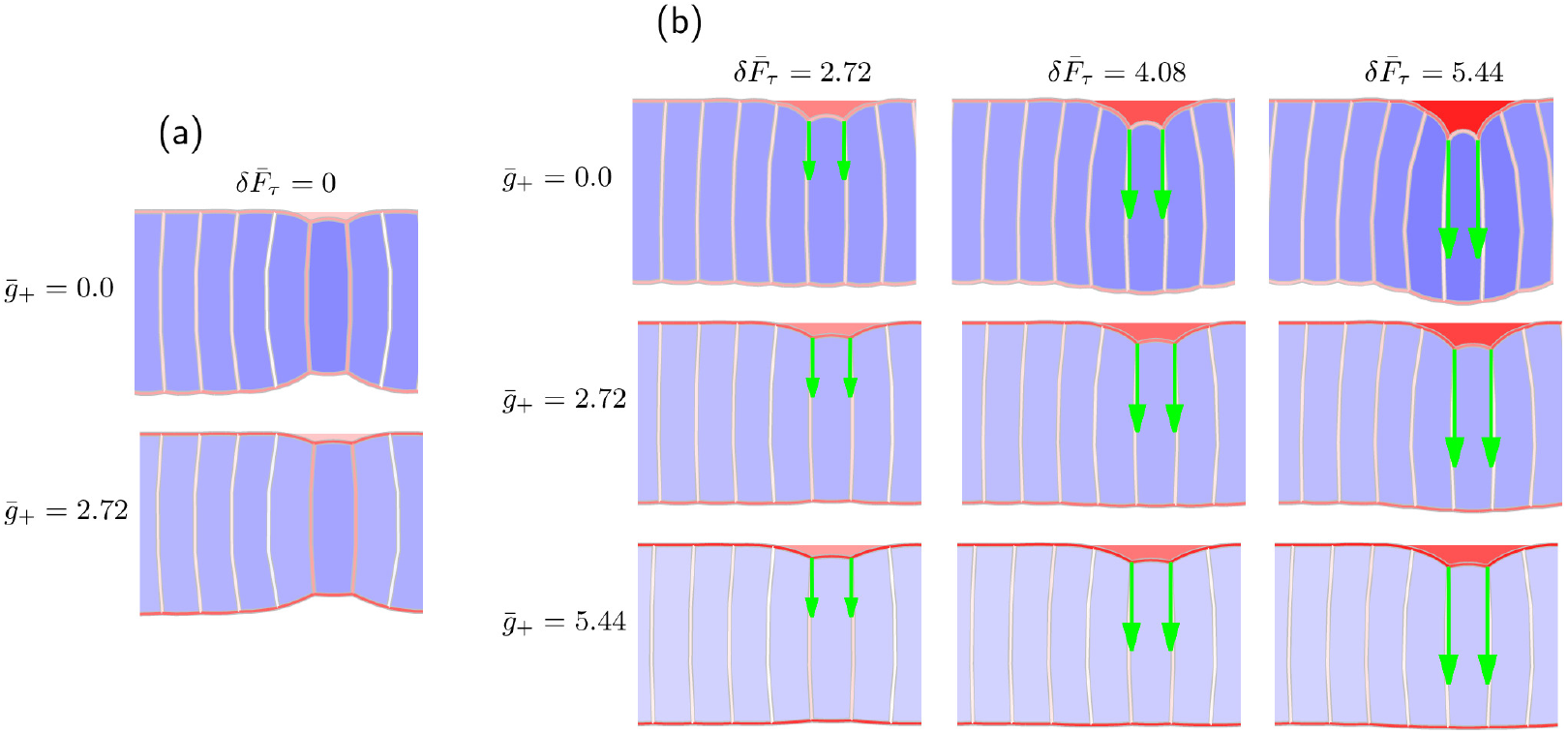
The effect of blastoderm tension 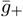 on the geometry of the invagination region during phase 1E of CFF. MNLVM results for (a) no inward force, 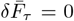, and (b) nonzero inward force, 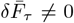. The values of 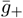 and 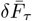, as labeled. The lateral membranes of the initiator cell are shortened by the factor 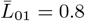 in panel (a) and by 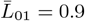 in panel (b).

Figure 9 illustrates the effect of apicobasal shortening of the initiator cell [Fig. 9(b)], of the tension-induced inward force (1) [Fig. 9(c)], and a combination of the shortening and inward force [Fig. 9(d)]. The inward force 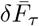 is exerted on the primary vertices at the junction of the apical and lateral membranes of the initiator cell, which is an approximate location of the actomyosin activity that produces tension along the future furrow cleft [31].

According to the simulation result shown in Fig. 9(b), the shortening of the lateral membranes of the initiator cell causes formation of the perivitelline fluid region above the initiator-cell apex, similar to the experimentally observed behavior depicted in Fig. 9(a). However, to maintain the force balance, an indentation is concurrently generated at the basal surface of the epithelial layer. Such a concave shape is not observed *in vivo*. Moreover, since the basal indentation is larger than the apical perivitelline fluid region, the center of mass of the cell goes up rather than down.

In contrast, when the inward force 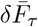 is applied, the perivitelline fluid region is generated without causing the indentation of the basal surface. Instead, the basal surface bulges out [Fig. 9(c)], but this artifact can be controlled by a moderate shortening of lateral membranes of the initiator cell [Fig. 9(d)]. Thus, the shape obtained in Fig. 9(d) matches the shape observed *in vivo*.

Experimental results indicate that during phase 1E of CFF, the apex of the initiator cell moves away from the vitelline membrane and stays below the blastoderm apices. The basal surface of the blastoderm remains flat [Figs. 4 and 9(a)], and an indentation is never observed in live embryos. Since this *in vivo* behavior cannot be reproduced *in silico* without the force 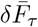, we conclude that the inward force associated with anisotropic tension developing along the furrow plays a critical role during the onset of the CF initiation process.

#### 2. Mechanical coordination of cell activities involved in CF initiation

It is interesting to ask how the coordination between the inward force 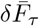 and initiator-cell shortening is achieved to obtain a well-formed CF initiation region. We propose that this is accomplished by mechanical feedback associated with the influence of the inward force on the lateral-membrane tension. Specifically, when 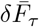 is applied, it causes a drop of the tension *σ*_l_ of the lateral membranes of the initiator cell [closed symbols in Fig. 10(b)]. We hypothesize that this tension reduction provides a mechanical signal that controls shortening of the lateral cortices.

This hypothesis is supported by a MNLVM simulation where the lateral tension is maintained on a constant level. The model shows that the constant tension condition

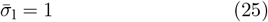

ensures that the basal surface remains nearly flat when the inward force 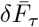 is increased [right panels of Fig. 10(a) and open symbols in Fig. 10(b)]. The resulting simulated configuration closely resembles the experimentally observed cell arrangement [Fig. 9(a)].

The above simulations indicate that the tension-induced inward force 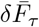 may play a dual role, i.e., it (*i*) mechanically drives the initiator cells away from the vitelline membrane (in the yolk direction) and (*ii*) provides a mechanical signal that helps synchronize cell activities. As demonstrated by Eritano *et al*. [31], the tension developing along the furrow also ensures that the activated initiator cells are well aligned. Thus, the tensile stress *τ* in the narrow strip along the furrow plays an important multifaceted role, which includes direct mechanical action in the inward and furrow directions and mechanical signaling.

#### 3. CF initiation in the presence of blastoderm tension

The simulations presented in Secs. V A 1 and V A 2 were performed for a system with no tension in the blastoderm. However, it is likely that *in vivo* such tension is present on both sides of the CF and plays an important role in mechanical signaling between morphogenetic movements [30, 47–49]. In Fig. 11 we explore the effect of normalized blastoderm-tension force 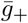 on the onset of CF initiation.

The results are shown for a system with no inward force 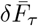 [Fig. 11(a)] and for several nonzero values of 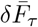 [Fig. 11(b)]. The simulations indicate that for a given degree of apicobasal shortening of the initiator cell and a fixed strength of the inward force, the tension 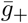 reduces the thickness of the perivitelline fluid region and increases its width. The force 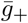 also causes a flat basal surface of blastoderm to slightly indent [Fig. 11(b) for 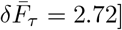 and protruding surface to flatten [Fig. 11(b) for 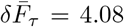 and 5.44]. In addition, the blastodermtension force 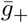 results in a reduction of cellular pressures (as indicated by the reduced intensity of blue in Fig. 11 for larger 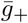) and a decrease of lateral-membrane tensions of the adjacent cells (the reduced intensity of red).

In summary, our results indicate that the onset of CFF entails a coordinated action of the inward force 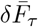 and shortening of lateral cell membranes of the initiator cells. The coordination is likely achieved via mechanical feedback associated with response of lateral-membrane tension to the inward force and the shortening. In the presence of blastoderm tension a stronger inward force is needed to produce a similar depth of the perivitelline fluid region, but phase 1E initiation can proceed in spite of the opposing tensile forces. In the following section we use MNLVM with a simplified pulley-point representation of the perivitelline fluid region (III B 3) to analyze the subsequent phases of CF initiation.

### B. Successive cells entering the cephalic furrow

This section focuses on CFF phases 1M/1L and 2E. (In our simulations we do not distinguish between phases 1M and 1L because cell nuclei are not represented in the model; see the nomenclature explained in Fig. 4). During phases 1M/1L and 2E, successive cells descend into the newly forming groove [Fig. 12(a)]. The descending cells undergo apicobasal shortening and basal expansion; concurrently with the expansion process, blastoderm progressively protrudes into the yolk sac. The cells at the edge of the protrusion narrow at the basal ends and expand near the apical ends to accommodate the changing orientation of the cells entering the furrow.

**FIG. 12.**
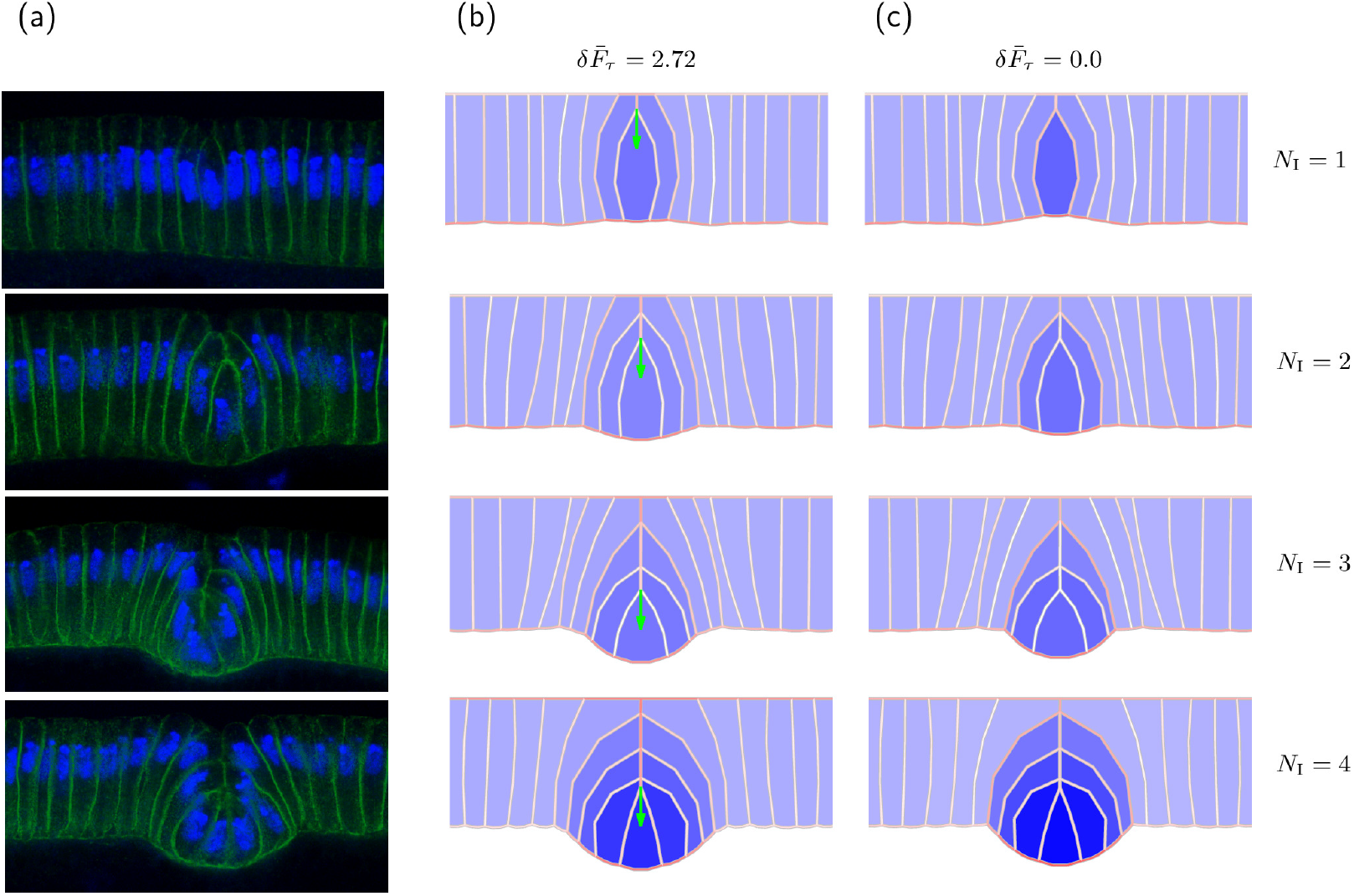
Invagination mechanics during phases 1M and 2E of CFF. (a) A time-lapse montage of fixed embryo images showing subsequent cells entering the CF groove. (b,c) Numerical reconstruction of the shapes shown in (a) for a system with tension-induced inward force 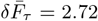 applied to the initiator cell apex [panels (b)] and no inward force, 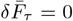 [panels (c)]. The inward force is indicated by the green arrows. The number of internalized cells *N*_I_ is indicated for each row.

Reproducing these shape changes in the vertex model requires precise numerical manipulation of the membrane and area parameters of multiple cells until the best match is obtained between the vertex system in mechanical equilibrium and the corresponding experimental image. To reduce the complexity of the simulation procedure, we use our simplified version of MNLVM, i.e., the variant with the perivitelline fluid region at the CF cleft modeled as a single pulley point [30], as described in Sec. III B 3 and Fig. 5(d).

Figures 12(b) and 12(c) show the simulated progression of CFF with and without the inward force 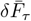, respectively. The simulation frames were obtained by adjusting system parameters until a good agreement was obtained between membrane lengths and curvatures of the cells in the simulation frames and the corresponding membrane features of the cells in the anterior (left) half of the experimental images.

In the consecutive simulation frames presented in Fig. 12(b,c), the initiator cell and *N*_I_ −1 cell pairs have already been internalized, i.e., descended below the blastoderm apices and are separated from the vitelline membrane by the subsequent cells in the invagination queue. The cells of the *N*_I_-th cell pair have met and their apical membranes have partially separated from the vitelline membrane. The experimental image in the row *N*_I_ = 1 corresponds to the phase 1M (flat basal surface) and the images in rows *N*_I_ = 2, 3, 4 correspond to the phase 2E (protruding basal surface).

The results depicted in Fig. 12 show that the significance of the inward force 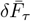 decreases when the invagination process progresses. The simulation images for *N*_I_ = 1 (phase 1M/1L) indicate that without the inward force 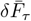, the basal surface is indented in the neighborhood of the initiator cell. Similar to the results discussed in Sec. V A 1, the force 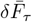 removes this indentation. When the basal membrane of the initiator cell expands during phase 2E (*N*_I_ = 2, 3, 4) the cell becomes asymmetric, and the pressure of the adjacent cells pushes it down. This mechanism allows the basal end of the initiator cell to be pushed below the blastoderm basis without the help of the inward force 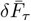.

The force 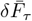 also helps form a well-organized furrow structure in the presence of blastoderm tension (Fig. 13). During phase 1M/1L (*N*_I_ = 1), the nonzero blastoderm tension 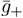 produces increased basal indentation, which is reduced by the inward force 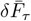 [Fig. 13(a)]. At the later stage of CF initiation (*N*_I_ = 4) tension force 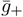 affects primarily non-invaginated cells at the edge of the newly forming CF protrusion, causing their shortening and generating inward curvature of the basal surface of the blastoderm [Fig. 13(b)].

**FIG. 13.**
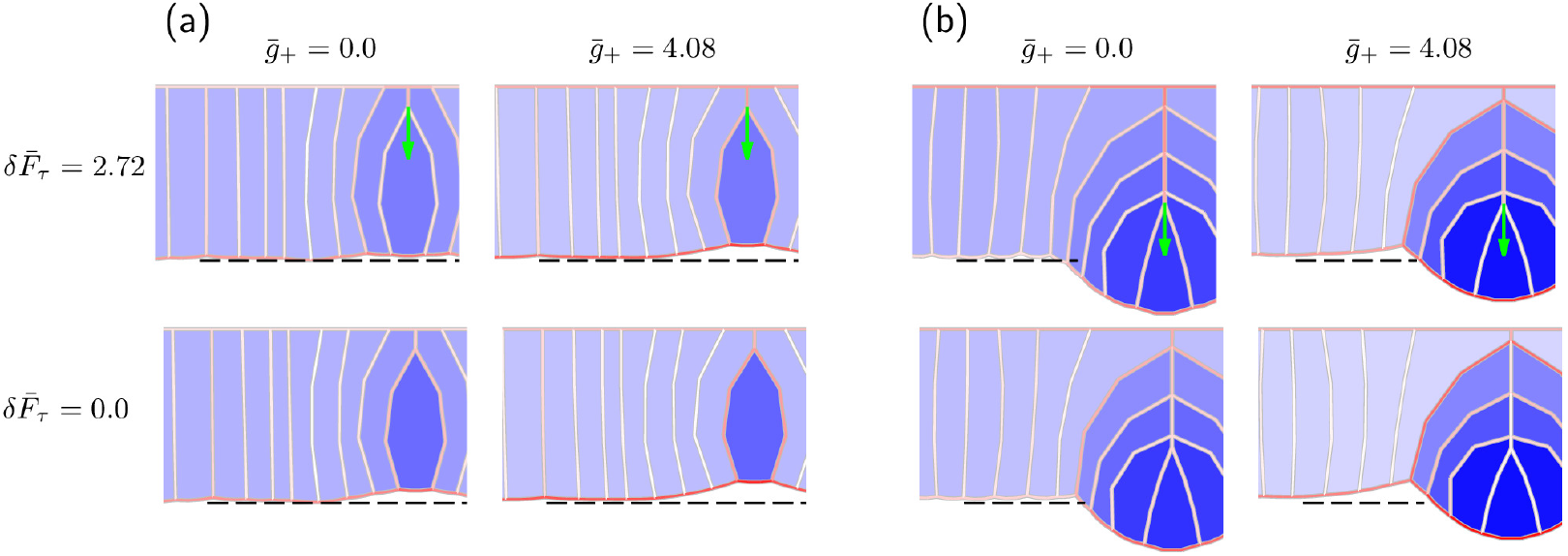
The effect of blastoderm tension on the geometry of the invagination region. MNLVM results for (a) *N*_I_ = 1 and (b) *N*_I_ = 4. The values of the blastoderm tension 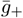 and inward force 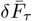 as labeled. The dashed line is the guide for the eye to highlight changes in CF geometry.

Since during phase 2E very similar cell configurations can be obtained with and without the inward force, we conclude that the force 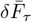 becomes redundant when phase 2E is reached, as far as the cross-sectional mechanics is concerned. However, since the underlying tension *τ* smoothly propagates along the furrow, the force 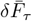, given by (1), helps coordinate the invagination progression between different cross-sections, smoothing out curvature differences and increasing CFF robustness. The evidence for this facilitating role of the tension-induced inward force can be found in the video S1 of the paper by Eritano *et al*. [31], which shows a coordinated line of initiator cells entering the furrow.

## VI. DISCUSSION AND CONCLUSIONS

Our *en face* force-center model, FCM, and multi-node lateral vertex model, MNLVM, taken together, provide a compelling picture of how cells in the developing embryo coordinate the initial stages of CFF. Our numerical simulations and analyses of experimental images (*i*) identify a combination of cellular-activity-driven local and longrange forces to produce the experimentally observed geometry of the developing furrow; (*ii*) shed light upon mechanisms that allow cells to collaboratively navigate the generation of tissue-scale changes by coordinating through mechanical-stress feedback. The strong agreement of our results with recent experimental findings [31, 46, 50–52] supports this mechanical-stress feedback-focused description of intercellular interaction.

Local forces that shape CF include membrane tensions and cell pressures. According to our present investigation and our previous study [30], while local stresses associated with membrane expansion or shortening and cellular pressures are sufficient to generate a sequence of cellular configurations observed during the later stages of CFF initiation (i.e., phases 1L, 2E, and 2L), we find that the initial phase (phase 1E), during which the initiator cell undergoes apicobasal shortening and its apex enters the furrow, cannot be produced by a combination of membrane tensions and cell pressures without an additional inward force directed towards the yolk sac.

Specifically, simulations using MNLVM show that apicobasal shortening of the initiator cells without the concurrent inward force produces a symmetric inward bowing of both the apical and basal surfaces. In contrast, *In vivo*, asymmetric generation of the perivitelline space paired with a flat basal surface is observed, and these shapes can only be produced with the help of an inward force that needed to ensure the force balance.

We propose that this required inward force results from the tension *τ* developing along the furrow. The existence of such tension was suggested by Eritano *et al*. [31] in the context of aligning initiator cells along the furrow cleft. Anisotropic tension along the furrow is likely generated by contractility of myosin II [see Fig. 14(a)], polarized along the furrow direction, with forces transmitted across adherens junctions.

**FIG. 14.**
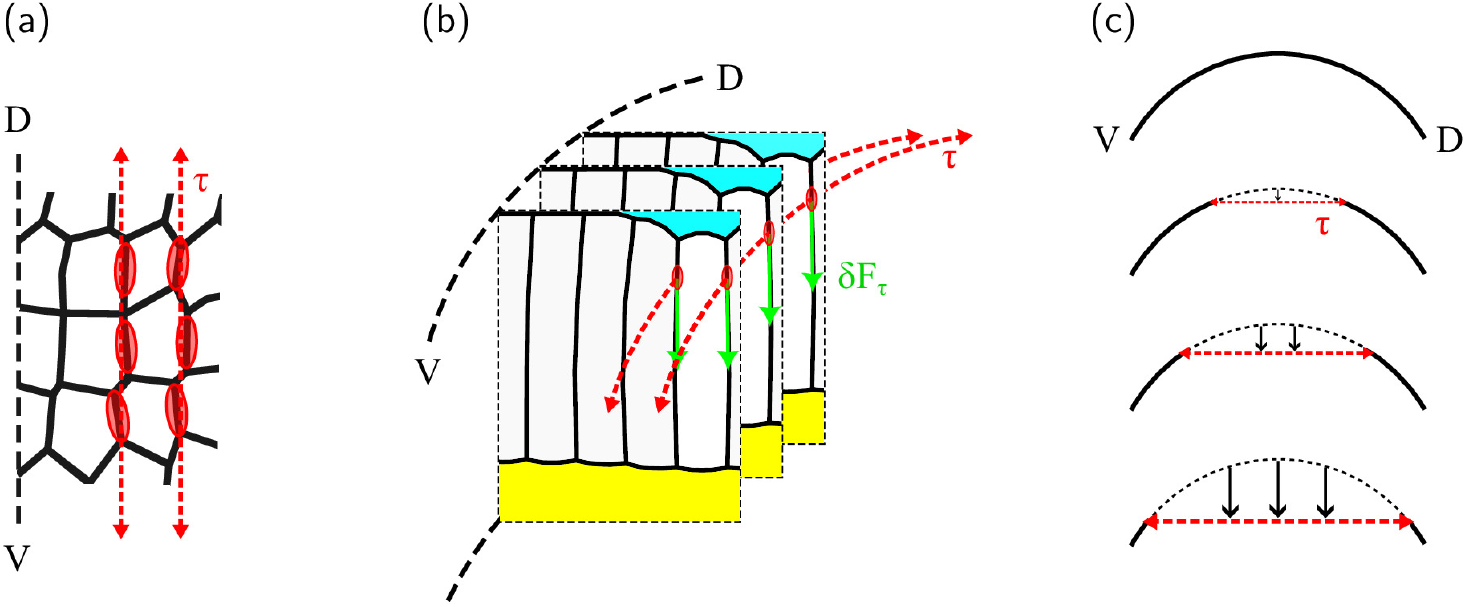
Tension along supracellular myosin II ribbons supports the tensile-stress-based activation of CF initiator cells, and further, is responsible for the initial descent of initiator cells into the furrow. The three panels illustrate our interpretation of experimental results from Eritano *et al*. [31]. (a) Eritano *et al*. [31] found that myosin is planar-polarized. Representations of myosin II accumulation are shown with red ovals; red arrows indicate tension directed along the dorsal-ventral (D-V) axis. Tension *τ* created by the myosin II ribbons along the curved epithelial layer generates an inward force *δF*_*τ*_ (green arrows). Perivitelline fluid region (cyan) and yolk region (yellow). Our simulation results show that both the tension-induced inward force and lateral shortening are required for proper cell conformation. We propose that the lateral shortening is controlled by mechanical-stress feedback and is a reaction to the drop in lateral membrane tension produced by the inward force *δF*_*τ*_. Increased tension *τ* along the supracellular myosin II ribbons produces increased inward force *δF*_*τ*_, which results in a gradual lowering (black arrows) of the furrow base (red dashed line).

The curved apical surface of the embryo allows the stress field to produce a tension-induced inward force *δF*_*τ*_ [Fig. 14(b)]. As demonstrated by our MNLVM, such a tension-induced inward force can provide the missing force contribution needed by the initiator cells to descend into the furrow.

Subsequently, this force progressively flattens the line of initiator cells while simultaneously forming and deepening the initial base of the furrow. [Fig. 14(c)] This description is consistent with the flattened line of invaginated initiator cells that can be observed in Video S1 in Eritano *et al*. [31].

Interestingly, the tension-induced inward force *δF*_*τ*_ appears to be indispensable only during the initial stages of CFF. Its presence during later stages seems to provide redundancy that increases the robustness of the overall process; the proper overall shape of the cell layer can, however, be achieved without the tension-induced inward force. Our results indicate that the presence of the inward force *δF*_*τ*_ at later stages, while not required, facilitates the progression of the furrow and ensures that CFF is a reliable and robust morphogenetic movement.

Based on our analysis of CF initiation patterns and stress propagation in a heterogeneous cellular system, we propose that furrow-scale tension *τ* not only mechanically sculpts the furrow structure in collaboration with local cellular forces, but it also provides mechanical signals that are crucial for coordination of cellular activity locally and across the entire furrow.

As demonstrated in our FCM, contractile activity within a given cell generates a mechanical stress field oriented along the furrow. This stress, which extends beyond the initiator cell’s immediate lateral neighbors, establishes a feedback mechanism whereby initiator cells are activated and, in turn, strengthen and further extend the stress field as myosin within their membranes constricts. Our FCM clearly shows that this kind of mechanical stress feedback mechanism leads to formation of constriction chains with transient discontinuities, similar to those observed *in vivo* (see Fig. 3).

The initiator-cell activation via a long-range tensile stress field likely contributes to the robustness of CFF. Neighbor-based initiator cell activation, similar to the one proposed by Popkova *et al*. [46], introduces the potential for a single cell along the forming furrow to be a critical point of failure. Similar to a domino chain, removal of one domino (in this case a disrupted activation of an initiator cell) can break the chain of activation and lead to a failure of CFF. In contrast, the inherent long-range propagation of the mechanical stress field allows relatively distant initiator cells to become activated, alleviating the possibility of individual cells becoming a critical point of failure.

The above mechanism promotes buildup of a coherent long-range stress field along the furrow. As suggested by our MNLVM, this stress not only produces the inward force *δF*_*τ*_ that pushes the initiator cells into the furrow, but also also provides mechanical signals to activate and guide initiator cell shortening. Specifically, the tension-generated inward force, acting along the lateral membrane, results in a reduction of the tension in the membrane below the point where the force is applied [see the schematic in Fig. 14(b)]. We have shown that when the membrane reacts to this reduction by restoring the initial tension, the correct overall shape of the cell layer is obtained.

Based on the above observations, we posit that CF initiation at a given position along the future furrow is not triggered by apicobasal shortening of initiator cells as (commonly assumed) but by the buildup of tensile stress propagating along the furrow. The associated inward force provides a mechanical signal that (*i*) triggers the initiator-cell shortening and (*ii*) controls its degree.

While our current investigations rely primarily on numerical simulations, knowledge of physical laws, and geometrical analysis of experimental images, we expect that the results presented here will inform future experimental and computational studies to further elucidate mechanical forces and feedback control governing CFF and other morphogenetic movements. In particular, further experimental studies of the underlying stress field and tension-induced inward force are necessary. Optogenetics could be used to selectively disrupt the mechanical activity of individual cells or groups of cells, providing a means to analyze the underlying mechanical stress field and to assess the reaction of affected cells to changes in the mechanical environment.

